# Genomic and phenotypic imprints of microbial domestication on cheese starter cultures

**DOI:** 10.1101/2024.03.19.585705

**Authors:** Vincent Somerville, Nadine Thierer, Remo S. Schmidt, Alexandra Roetschi, Lauriane Braillard, Monika Haueter, Hélène Berthoud, Noam Shani, Ueli von Ah, Florent Mazel, Philipp Engel

**Affiliations:** DMF, University of Lausanne, Switzerland; Agroscope, Switzerland; Université Laval, Canada; McGill, Canada

## Abstract

Domestication – the artificial selection of wild species to obtain variants with traits of human interest– was integral to the rise of civilization. During the neolithic, the oversupply of food enabled by the domestication of crop plants and farm animals was associated with the invention of food preservation strategies through microbial fermentation. However, it remains unclear whether fermented food microbes show similar signs of domestication by humans like plants or animals. Only a few eukaroytic have been studied so far in this respect (e.g., yeasts used in mantou or wine), whereas little is known for bacteria.

Here, we tested if cheese starter cultures harbour typical hallmarks of domestication by characterising over 100 community samples and over 100 individual strains isolated from historical and modern traditional Swiss cheese starter cultures. We find that cheese starter cultures have low genetic diversity both at the species and strain-level and are taxonomically and phenotypically stable. Our analyses further suggest that the evolutionary origin of the bacteria in cheese starter cultures coincided with the start of cheesemaking as reported from archeological records. Finally, we find evidence for ongoing genome decay and pseudogenization via transposon insertion related to a reduction of their niche breadth.

These characteristics suggest that cheese starter cultures were domesticated by humans before knowing about microbes, potentially starting as early as the neolithics Future work documenting the prevalence of these hallmarks across diverse fermented food systems and geographic regions will be key to unveiling the joint history of humanity with fermented food microbes.

## Introduction

Domestication is the process of modifying wild species through artificial selection to the benefit of a “domesticator”, which is usually human (1,2). This process was integral to the rise of human civilization (3–5). In particular, domestication of both crop plants (6) and farm animals (7) during the neolithic agricultural revolution around 1-10 millennia B.C. (4) is a cornerstone of human history that allowed in the long run the emergence of non-food producing sectors (e.g. science and philosophy) and initiated large-scale anthropogenic changes of the earth biosphere (8). Plant and animal domestication is characterised by tremendous phenotypic and genetic changes from the wild ancestor, such as an increase in body mass (9) or change in nutrient content (10) which overall contributed to the production of food surplus.

During the neolithic, this oversupply of food was associated with the invention of food preservation strategies by microbial fermentation – a metabolic process that converts sugars into acids – to produce, for example, fermented vegetables, wine or cheese (11,12). This leads to a decrease in pH, which reduces undesired microbial growth and prevents spoilage of stored food. Subsequently, fermented food products have diversified in a myriad of forms all over the globe (13,14) constituting healthy and tasty components of the diet which are key in many cultures and sustainable opportunities for the future of food in others (15). This raises the fundamental question whether, as for plants and animals, fermented food microbes have also been domesticated by humans, and if so when and how did this happen.

It is commonly accepted that some fermented food microbes have been maintained through continuous passaging (16,17) and artificial selection for specific traits (e.g. shelf life or taste). Accordingly, we expect them to present typical “hallmarks” of domestication, i.e. genomic and phenotypic signatures associated with (microbial) domestication that distinguish them from their wild counterparts. However, only a handful of microbial domestication cases have been documented empirically (18), mostly for eukaryotic microorganisms. For example, Saccharomyces cerevisiae used for bread making (19) and alcoholic fermentation (20,21) or *Aspergillus oryzae* used for sake, soy sauce, and miso production (22) show genomic and phenotypic characteristics that distinguish them from their wild counterparts (23,24).

Thus, while the genomic signatures of domestication are well defined for microbial eukaryotes, we know relatively little about the collective genomic and phenotypic consequences of bacterial domestication used for food fermentation. Also, it remains unclear whether and when bacteria have been domesticated (24,25). For example, *Oenococcus oeni*, which is responsible for the malolactic conversion in winemaking, is thought to have rapidly diverged from its ancestor due to the emergence of hypermutator strains, but it remains unclear to what extent subsequent evolution was influenced by human domestication (26). Similarly, strains of *Lacticaseibacillus paracasei* isolated from ripening cheese show signs of adaptation to milk but lack other hallmarks of domestication (27). Altogether, this suggests that most suspected cases of domestication in fermented food bacteria lack a complete characterisation of domestication hallmarks previously observed in microbial eukaryotes like yeast.

To explain this discrepancy, we posit that, in contrast to yeasts (i.e. in beer, bread and mantou), the previously studied bacteria have not been continuously passaged with rapid and iterative bursts of growth solely on the fermented foods but rather persisted in food-associated environments (e.g. grape skin for *Oenococcus oeni*). This reduces the strength with which artificial selection can act on evolution. In contrast, thermophilic cheese starter cultures are a promising candidate to test for domestication in bacterial communities used for fermentation because they have been passaged and selected for thousands of years via backslopping (i.e. continuous re-inoculation of previous-day whey) and have been extensively selected for flavour and rapid acidification purposes (28). Thermophilic starter cultures are used at high (above 42°C) temperatures and are dominated by three thermophilic bacteria, namely *Streptococcus thermophilus*, *Lactobacillus delbrueckii* subsp. *lactis* (hereafter only *L. delbrueckii*) and *Lactobacillus helveticus* (*29*). While analyses of a limited number of genomes from these species have revealed signs of genome decay (18,30–34), we currently lack a systematic definition and screening of domestication hallmarks on these microbial communities (16,17).

In this study, we aimed to detect signs of domestication in thermophilic cheese starter cultures and to date the potential domestication events. To this end, we collected both modern and historic (1970s) samples of 11 undefined cheese starter cultures that are continuously passaged as undefined cheese starter cultures (described in (35)) and used to make three cheese varieties in different regions in Switzerland (Fig. 1A). We characterized over 1000 samples phenotypically, and about 100 metagenomes and more than 100 bacterial isolates from historical and modern cheese starter cultures genetically (Fig. 1B). Moreover, by carrying out experimental evolution assays with the major community members we expanded on previously proposed hallmarks of domestication for eukaryotes to define five specific hallmarks of microbial community domestication for bacteria (24): i) phenotypic stability over time, as a result of the selection for food preservation, ii) simple and stable microbial diversity both at the species and strain-level, as a result of continuous passaging, iii) evolutionary origin of focal species coinciding with the start of food preservation, iv) gradual genome decay and v) adaptation to the food environment by the gradual reduction of niche breadth. Collectively, these results suggest that thermophilic cheese starter cultures have been domesticated by humans for millennia.

**Figure 1.**
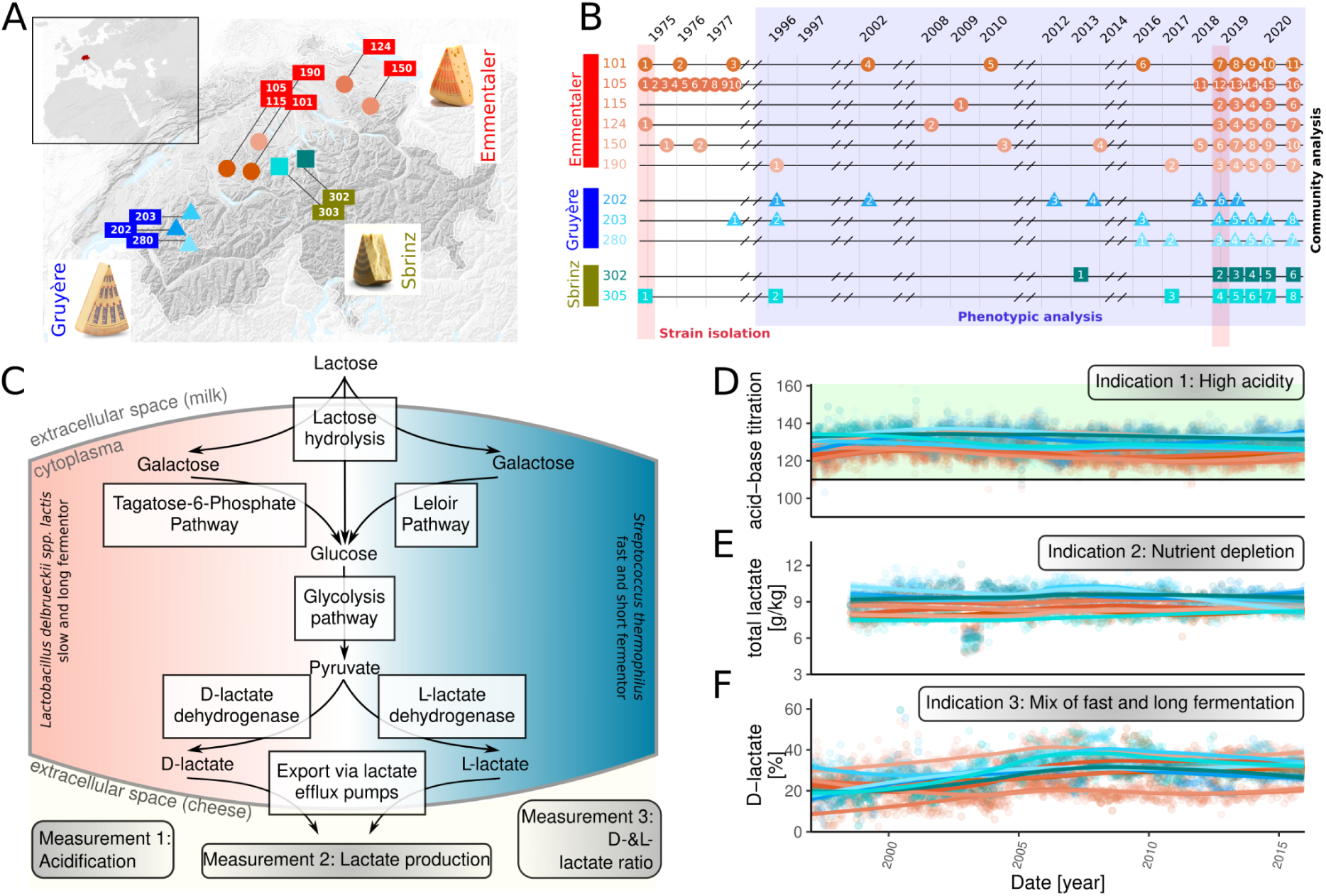
Phenotypic stability of thermophilic cheese starter over 50 years of cheese production. A) Origin of 11 continuously passaged starter cultures originating from different regions in Switzerland and different cheese varieties (the inlet shows the location of Switzerland within Eurasia.) B) Both modern and historic sampling time points for the community analysis (colored circles, triangles, and squares), phenotypic analysis (blue background) and strain isolation (red background) of all starter cultures. C) The general mechanism of food preservation for thermophilic starter cultures consisting of a stable i) acidification, ii) lactate production and ii) D- &L-lactate ratio. Phenotypic measurements of the starter cultures D) total acid-base titration value, E) total lactate and F) percent D-lactate in respect to L-lactate.

## Results

### #1. Food preservation by stable acidification as a result of lactose fermentation by thermophilic cheese starter cultures

The first hallmark of domestication predicts phenotypic stability over time, as a result of continuous passaging in a stable environment (i.e. milk) and selection for a specific trait, i.e. food preservation, as determined by a charcteristic smell and favorable process parameters. The underlying phenotypic properties that give rise to enhancing the shelf life of dairy products by preventing the growth of undesired microbes are i) the acidification of the environment and ii) the reduction of the amount of easily available nutrients. This is accomplished by rapidly and reliably transforming lactose into lactate (homofermentation), notably consisting of the following five steps: i) lactose hydrolyzation, ii) galactose transformation either by the Leloir- or the tagatose-6-phosphate pathway, iii) glycolysis, iv) L- and D-lactate production by *S. thermophilus* and *L. delbrueckii*, respectively and v) lactate export by an efflux pump (36–40) (Fig. 1C).

In the following, we measured the three key phenotypes for food preservation by lactose fermentation, namely acidification, lactate production, and D-&L-lactate ratio (Fig. 1C). Firstly, we titrated >1000 samples, and found that acidification potential was consistently high (>110 ml of 0.1 M NaOH, which is generally regarded as good acidification in cheese production) and stable over time (Fig. 1D, mixed effect linear model time slope estimates with p<.05). Secondly, the amount of detected lactate and the ratio of the corresponding enantiomers, D- and L-lactate, slightly but not dramatically increased over time (Fig. 1E/F, mixed effect linear model time slope estimates with p<.05). The former suggests a stable nutrient depletion by accumulation of lactate as a final product. The latter suggests that the relative metabolic contribution of the two dominant community members is similar across time. This is essential as *S. thermophilus* ferments rapidly but only until pH 5, whereas *L. delbrueckii* commonly starts significantly fermenting from pH 5 downwards for a longer time. In summary, these results show that cheese starter cultures are highly stable in their phenotypic properties, as expected when communities are selected for specific traits.

### #2. Cheese starter cultures are simple and stable microbial communities

The second hallmark of domestication predicts that the microbial diversity, both at the species and strain-level is simple and stable as a result of the passaging in a contained and highly stable nutrient-rich environment. We tested this by selecting a total of 98 cheese starter cultures for shotgun sequencing (6-11 mio. reads per sample, circles, triangles, and squares in Fig 1B) and determined their taxonomic composition using a short read taxonomic profiler (mOTU2, (41)). As expected, we found that most samples were dominated by only two species, *Streptococcus thermophilus* and *Lactobacillus delbrueckii* subsp. *lactis* (Fig. 2A). Yet, we noted two apparent signs of instability through time: (i) *L. helveticus* was only present in early samples of some cheese starters and (ii) the relative abundance of *S. thermophilus* and *L. delbrueckii* varied across samples of the same starter culture (Fig. 2A). The former observation suggests that *L. helveticus* may have been lost over time without changing the phenotypic properties of the starter cultures. The latter observation may be due to the fact that the precise sampling timepoints of the starter cultures were not controlled for and that *L. delbrueckii* growth is delayed relative to *S. thermophilus* (*35*). Altogether, this suggests that the species-level composition of the cheese starter culture was remarkably stable over nearly 50 years of sampling.

**Figure 2.**
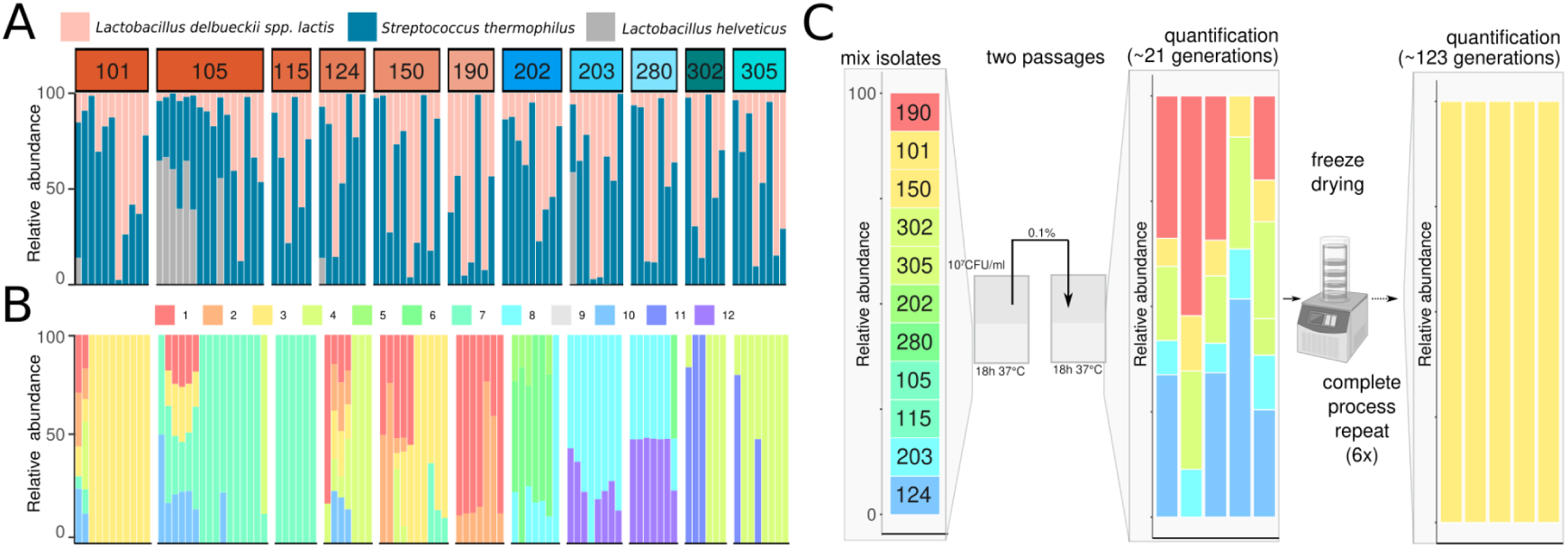
Simple and stable community composition both at the species and strain-level in cheese starter cultures. A) Species-level composition across all samples: within each starter culture, samples are ordered from older to more recent. B) Strain-level compositional diversity across the different samples for the 12 different *S. thermophilus* strains identified (each strain is representative of a sub-species clade). C) The propagation experiment of all dominant *S. thermophilus* and *L. delbruecki* (*L. delbrueckii* is not illustrated as it only included one strain) over ∼21 and ∼123 passages.

To assess the within-species diversity, we mapped the metagenomic reads against a reference database containing isolate genomes of each of the three species and quantified the number of polymorphic sites detected in core genes (i.e. genes identified in all strains of a given species, see methods) in each sample. The proportion of polymorphic sites was similar over the samples with around 0.11 % (SD=0.72 %), and 0.02 % (SD=0.03 %) for *S. thermophilus* and *L. delbrueckii,* respectively (SFig. 1), which is comparatively low with respect to bacteria found in non-food fermentation systems like in the gut of animals (3 % and 2-10 % polymorphic sites within species in the gut microbiota of human (42) and honey bees (43), respectively).

To determine the actual number of strains the detected within-species diversity corresponds to, we genotyped > 2000 colonies from the 11 cheese starter cultures (Fig. 1B) and sequenced the genomes of 112 isolates. Using an all-against-all genomic distance analysis implemented in poppunk (44), we found that the sequenced genomes separate into 12 *S. thermophilus*, two *L. delbrueckii* and two *L. helveticus* sub-species clades (see methods). Overall, the sub-species clades within the different species are very similar (min ANI: Sterm=98.6 %, Ldel=98.9 %, SFig. 2 (45)). These sub-species clades accounted for most of the SNPs detected by metagenomic sequencing (93% and 78% of the metagenomic SNPs from *S. thermophilus* and *L. delbrueckii,* respectively, Fig. 2B, SFig. 3), suggesting that we have isolated and sequenced most of the sub-species diversity present in the analyzed cheese starter cultures. In the case of *L. delbrueckii* and *L. helveticus*, a single sub-species clade dominated in all analyzed samples, while for *S. thermophilus*, one to four sub-species clades per sample were detected.

**Figure 3.**
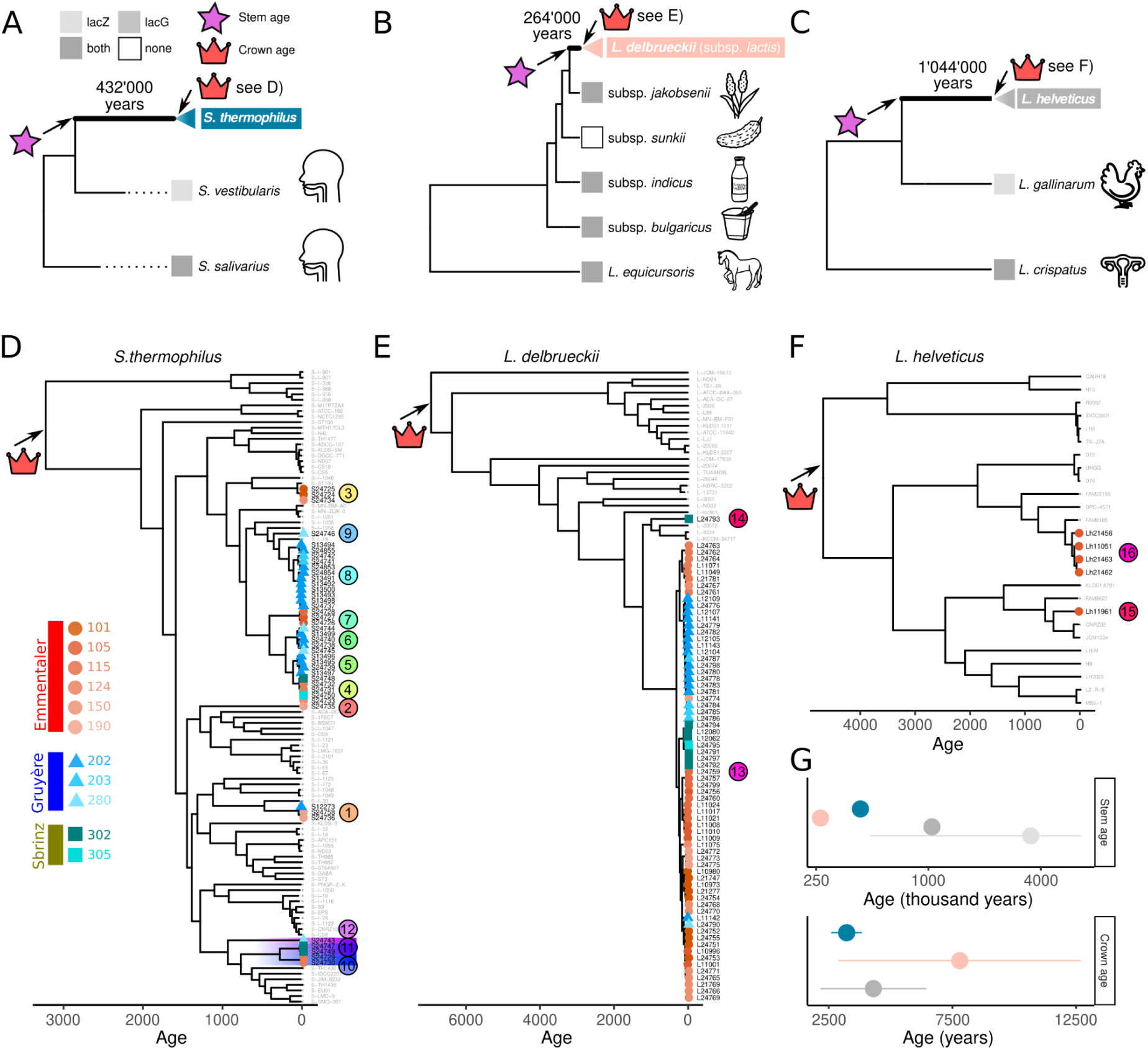
Phylogenetic dating of the origin of cheese-starter bacteria. A)B)C) Maximum likelihood rRNA operon (∼5kb) phylogeny of the focal species and their closest sister (sub)species. Tips are annotated with labels indicating 1) the presence of lacZ or lacG genes within the genomes and 2) the sampling origin of the different isolates (depicted by silhouettes). Stem age is depicted on the phylogeny with a star and annotated with its estimated age, along with a representation of sub-species diversity, depicted with a triangle that represents the phylogeny of panels D)E)F). D)E)F) Strain-level phylogeny illustrates recent diversification of cheese starter sub-species clades for D) *S. thermophilus*, E) *L. delbrueckii* and F) *L. helveticus.* Different sub-species clades are highlighted in colour. G) The stem age is calculated by the molecular clock of the rRNA operon for the different species and genera (also annotated near the stem nodes in panels A)B)C). The cheese starter culture-related species are highlighted with the corresponding colors (*L. delbrueckii*=pink, *L. helveticus*= grey and *S. thermophilus*=blue). The light grey point illustrates the stem age of all other species in the lactic acid bacteria family. The crown ages are inferred by a molecular clock rate based on the core genome phylogeny and historical samples as calibration points after removing putative horizontal transfer events.

Finally, to test if the milk environment in which these bacteria have been passaged cannot carry a larger amount of diversity at the sub-species level, we combined one randomly chosen *L. delbrueckii* (all from the same sub-species clade) and one *S. thermophilus* isolate from each cheese starter culture at equal amounts and co-cultured them in five replicates for 27 passages (∼123 generations) (Fig. 2C). After 21 generations, we found that five of the 11 original *S. thermophilus* strains were still present. However, after 123 generations, only one strain could be detected in the community (Fig. 2C).

In summary, these results show that undefined cheese starter cultures have a simple community composition at the species and strain-level that shows a high degree of stability in partuclar at the species level likely due to the continuous passaging in a stable, closed and nutrient-rich environment.

### #3. The onset of cheese starter strain diversification coincides with the origin of dairy fermentation in humans

The third hallmark of domestication predicts that the evolutionary origin of cheese starter bacteria should coincide with the start of cheesemaking or dairy fermentation as reported from the archeological record about 7000 years ago in Europe (11). To do so, we sought to date the evolutionary transition(s) from non-dairy environment to dairy environment using phylogenetic comparative methods. This approach first maps strain or species habitat preference (dairy/non-dairy) onto a phylogeny to identify the evolutionary transition(s) between habitats and then, in a second step, uses the molecular clock and the dates of historical samples as calibration points to estimate the age of this transition.

To identify the evolutionary transition(s) from non-dairy to dairy environment. We assembled a genomic database of 234 strains from the three cheese starter bacteria as well as closely related species by combining our own dataset with publicly available data (complete list of genomes provided in Supp. table 1). We mapped the isolation source on whole genome phylogenies (for within-species phylogenetic relationships) and on rRNA phylogenies (for between-species relationships). Most strains from *S. thermophilus, L. delbrueckii* and *L. helveticus* were isolated from cheese-starter culture or other milk products with a few isolates from fecal samples probably originating from a dairy diet (SFig. 4). Corroborating previous findings (46), ancestral niche reconstruction suggests that with a high likelihood of 99.99%, 99.99% and 76% the niche of the ancestors of all known strains of *S. thermophilus, L. delbrueckii* subsp. *lactis* and *L. helveticus* was already associated with dairy products (Maximum likelihood model, SFig. 5). In addition, although the three closest related sister species to our focal species are of animal rather than dairy origin, they all encode enzymes to degrade milk (LacG or LacZ galactosidases, Fig. 3A-C). This suggests that these milk-adapted sister clades probably evolved with the appearance of milk in mammals roughly 200 mio. years ago (47) but that only our focal species are found in the dairy environment.

**Figure 5.**
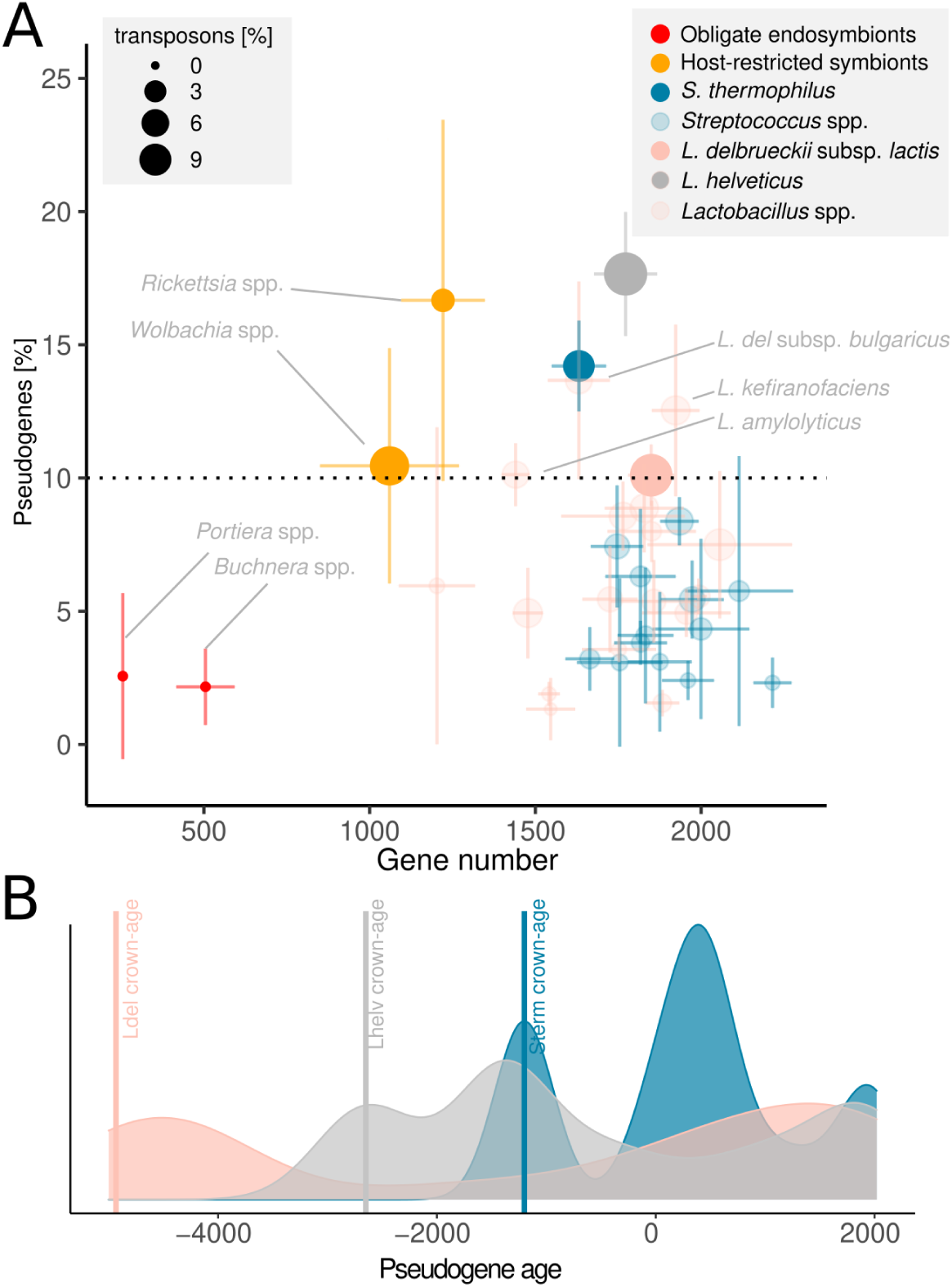
Dating of ongoing genome decay by pseudogenization. A) The fraction of pseudogenes per genome vs. the gene count in the genomes of two obligate endosymbionts (*Buchnera* spp. and *Portiera* spp.), and host-restricted symbionts (*Wolbachia* spp. and *Rickettsia* spp.), the three focal species (*S. thermophilus, L. delbrueckii* and *L. helveticus*), and two closely related species *(L. amylolyticus, L. kefiranofaciens*). The size of the points indicates the percent of transposons detected in the genomes; the error bar indicates the standard deviation. *S. thermophilus*: mean=1657 genes, std=47; *L. delbrueckii*: mean=1868 genes, std=33 and *L. helveticus*=1772, sd=95. % of pseudogenes: *S. thermophilus*: mean=14%, std=1%; *L. delbrueckii*: mean=10%, std=0.6% and *L. helveticus*=18, std=2.3,) B) The smoothed density plot of the approximate date of the pseudogenes is estimated from the most recent common ancestor of all taxa containing a given pseudogene. Vertical lines indicate the MRCA of all sequenced strain within a species.

Therefore, we propose that the origin of cheese starter bacteria is located somewhere between the split from the sister species (stem age) and the most recent common ancestor (MRCA) of the strains within the species (crown age) of the three cheese starter culture species (branches highlighted in black in Fig. 3A-C).

To estimate the age of the non-dairy to dairy transition, we took advantage of two independent molecular clocks of the stem and crown age. Stem ages were determined by using the divergence time from the rRNA phylogeny. Assuming an rRNA substitution rate of 1 substitutions/site per 100 million years (48), we estimated the divergence of *S. thermophilus*, *L. delbrueckii* subsp*. lactis* and *L. helveticus* from their sister taxon at around 432’000, 264’000 and 1’044’000 years ago, respectively (Fig. 3A-C). While this is more recent than most other species within the two genera (Fig. 3G), it is still 3 order of magnitudes older than the first report of dairy fermentation (7000 years ago). Crown ages were determined using dated core-genome phylogenies of our focal species (Fig. 3D-F, SFig. 6). Using a molecular clock based on the core genome phylogeny and historical samples as calibration points (49), we estimated a substitution rate of 1.1 SNPs per clonal core genome and year, which falls in the average range typically observed for other bacterial species (50,51). We extrapolate that the crown age of *S. thermophilus*, *L. delbrueckii* subsp. *lactis* and *L. helveticus* is around 3221, 7798 and 4304 years ago, respectively (Fig. 3G). This indicates that the known strain-level diversity within species may have started to emerge within the neolithic, overlapping the origin of dairy fermentation in Europe about 7000 years ago (11).

**Figure 6.**
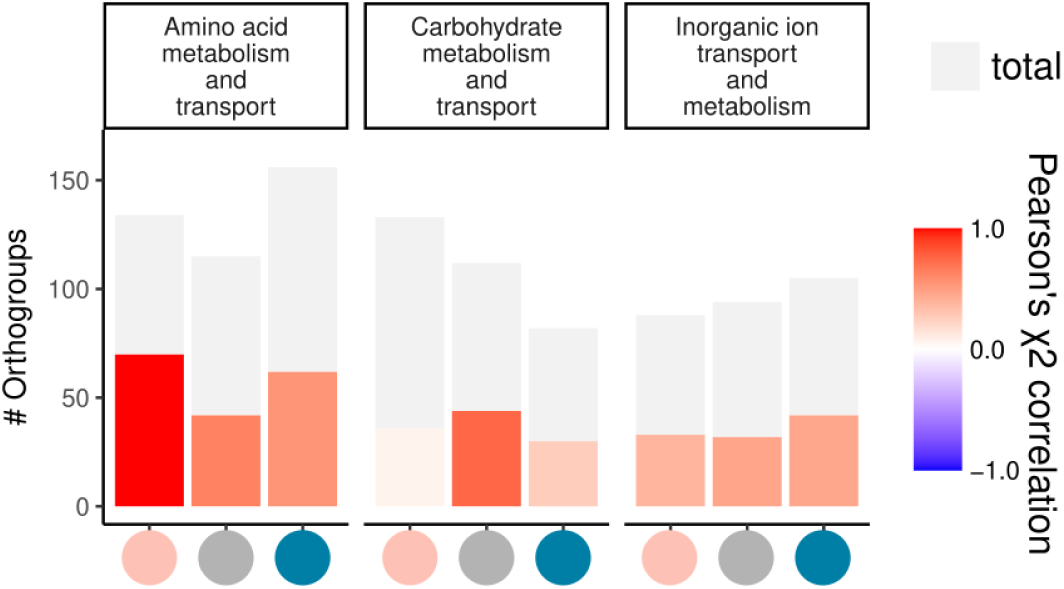
Enriched functions in pseudogenes. The total number of orthogroups (grey) and the Orthogroups containing pseudogenes colored if overrepresented (red) based on the Pearson Chi-square test (scale indicated in the figure). The three different species (*L. delbrueckii*=pink, *L. helveticus*= grey and *S. thermophilus*=blue) are separated accordingly by color.

To further elucidate the mode of subsequent diversification of strains, specifically from the cheese starter cultures across different geographical locations and cheese varieties, we mapped the isolation source to the strain phylogenies (Fig. 3D-F, SFig. 7). We found that, while there are exceptions, cheese varieties generally clustered on the phylogenies, i.e. bacteria isolated from different cheeses usually form distinct monophyletic groups. This suggests that environmental (different milk or cheese preparation techniques) or dispersal limitations constrained the subsequent diversification of cheese starter cultures.

Overall, this indicates that while the split from the next closely related known species is many hundred thousand years ago (stem age, Fig. 1D/E), the known strain-level diversity within the species (i.e. crown age) has likely started to emerge within the neolithic (<10’000 B.C.) coinciding with the start of dairy fermentation.

### #4. Cheese starter bacteria show genome decay by transposon expansion

The fourth hallmark of domestication predicts that cheese starter bacteria show signs of gene loss and genome decay (52). This is expected when bacteria thrive in stable and recurring environments with extensive nutrient availability and relatively small population sizes (53). This hallmark additionally predicts that the onset of genome decay must have started after the onset of domestication itself. To test this hallmark, we compared the genome size and degree of transposon insertion of our focal cheese starter strains to their closest relative. To put these estimates into a broader context, we also included data on insect-associated bacterial endosymbionts that shows extensive genome decay due to pseudogenization (54).

All three cheese starter culture species had smaller genome sizes than those of closely related species (Wilcoxon test, p<0.05, Fig. 5A, SFig. 8) and a higher number of pseudogenes (Wilcoxon, p<0.05, Fig. 5A, SFig. 9). We found that many of the detected pseudogenes in the three cheese starter culture species were the result of insertion events of transposons (mean=45%, std=9%; SFig. 10) belonging to 15 different IS element families (SFig. 11). Three other *Lactobacillus* spp. had similar signs of genome decay, all of which were also food-fermentation-associated species, namely *L. delbrueckii* subsp. *bulgaricus*, *L. kefiranofaciens* and *L. amylolyticus*. The observed genome decay of our focal genomes is in the range of host-restricted symbionts such as *Rickettsia* spp. and *Wolbachia spp.*, but not as extreme as for obligate endosymbionts of plant-sap feeding insects that have co-evolved with their host for millions of years. Overall, our findings suggest that all three cheese starter culture species have experienced pseudogenization via transposon insertion during their evolutionary history.

If genome decay started at the onset of domestication as predicted from the hallmark, the timing of pseudogenization events should be more recent than the onset of domestication itself in the neolithic. To estimate the onset of genome decay, we reconstructed the evolutionary history of pseudogenization events by mapping the presence/absence of modern pseudogenes onto the strain phylogenies. For each pseudogene, we identified the most recent common ancestor (MRCA) of the genomes which contained the pseudogene or in which the gene was completely missing (assuming the pseudogene was lost, SFig. 12). We found that most pseudogenes likely originated in the last millennia (Fig. 5B). The overall age similarity between the loss of functional genes (i.e. pseudogenization) and the cheesemaking development (within the last couple millennia) suggests that the persistent bottleneck and selection pressures of cheesemaking shaped the genomic evolution of these species. The ∼ 3000 years lag between the start of the cheese making and the estimated start of pseudogenization could arise if cheese making remained a spontaneous fermentation process for millennia before being more tightly controlled by backslopping. We conclude that the three focal species show clear signs of recent and ongoing genome decay.

### #5. Reduction of the niche breadth and adaptation to the cheese making environment

The fifth hallmark of domestication predicts a reduction of the niche breadth associated to pseudogenization of non-essential genes driven by adaptation to the cheese environment. We first tested the ability of cheese starter culture bacteria to grow on a wide range of carbon sources that are representative of diverse non-dairy environments. We found that *S. thermophilus* and *L. delbrueckii* can only metabolize respectively 5 and 9 of the 92 tested carbon sources, while their closest non-dairy relatives could metabolize respectively 12 (58% drop) and 61 (85% drop, SFig. 13-14).

We then sought to explore whether the pseudogenization of genes observed in these species could explain their specialization to the cheese environment and the few carbon sources used. To do so, we tested whether cellular functions - in particular carbon metabolism - were enriched in pseudogenes. Pseudogenes were spread across many orthologous gene families (OGs): from all 19,728 OGs in all three species, we identified 5,639 (29%) containing at least one pseudogene. Strikingly, three COG categories were overrepresented among the OGs containing pseudogenes for the three cheese starter culture species independently: carbohydrate (G), amino acid (E) and inorganic ion transport and metabolism (I) (χ2>0.3, p-value<0.05, Fig. 6, SFig. 15). More specifically, the three KEGG metabolic modules i) pentose phosphate pathway (ko00030), ii) fructose and mannose metabolism (ko00051) and iii) starch and sucrose metabolism (ko00500) were commonly pseudogenized (SFig. 16).

To test if the identified pseudogenes are indeed not or less expressed, we carried out RNA-seq of cheese during the first 24 hours of cheese making. We observed that pseudogenized genes are generally less expressed than non-pseudogenized genes throughout the first 24 hours of fermentation (RNAseq experiment, SFig. 17), suggesting that in general they are metabolically less important.

In summary, these results show that recent pseudogenization affected genes involved in the degradation of carbohydrates that occur in plants but not in milk, suggesting that cheese-making strains have lost genes that appear non-essential today in a stable and predictable dairy environment but were likely more important in their ancestral niche.

## Conclusion

Overall, fermented foods represent a stable, microbe-rich environment where human-controlled continuous passaging and selection results in the establishment of defined and stable microbial communities with specific phenotypic properties. So far, apart from the domestication of some eukaryotic microbes (e.g. Saccharomyces cerevisiae used in beer, wine or mantou), we know surprisingly little about the history of fermented food microbes, and whether they have been domesticated as extensively as cattle and crop plants remains an open question (24,25). We can conclude from our study that starter cultures show clear signs of domestication. i) They are highly stable in their acidification and lactose utilization, likely because of ongoing selection of the preservation properties (hallmark 1). ii) They contain simple and stable microbial communities maybe because of continuous passaging (hallmark 2). iii) The origin of the strains can be dated back to the likely emergence of dairy fermentation and the resulting anthropogenic selection pressure (hallmark 3). iv) They show clear signs of recent and ongoing genome decay that can be expected from the stable and nutrient-rich environments with relatively small population sizes and continuous passaging (hallmark 4). v) They show a reduction of the niche breadth and adaptation to the cheese making environment suggesting that their current niche is restricted to dairy fermentation batches (hallmark 5). We acknowledge that this might not be a complete list of domestication hallmarks. Most notably, recent studies have looked at the evolution of the pangenome of the different clades of *L. delbrueckii* (*34,55*) and *S. thermophilus* (*33*). While these studies similarly conclude that these species have a distinct evolutionary history tied to dairy fermentation, it currently remains unclear if the pangenome is open and expanding (33). Notably, the accessory genome contains numerous carbohydrate utilization genes suggesting a broad nutrient range. Here, we suggest that the size of the pangenome is likely also a consequence of extensive and ongoing pseudogenization because of a substantial fraction of highly mobile genes in the accessory genome namely active transposases, phages or phage defense genes (34,35,56,57). In any case, the addition of more high-quality genomes and the broader sampling in dairy and non-dairy environments will enable us to understand the pangenome diversity better and explore how it relates to domestication history. Collectively, we suggest that thermophilic cheese starter cultures have been domesticated by humans for millennia, probably starting during the neolithic when cheese-making emerged. While the exact timing of the transition from a wild dairy fermentation to backslopping is not entirely known, the bacteria have likely been in repeated cycles of selection for rapid and reliable acidification and also genetic drift through the repeat subsampling by backslopping. Our study fills an important knowledge gap by addressing if, when and how microbes have evolved to the anthropogenic usage and provides a conceptual framework that can be applied to other fermented food products. For example, from the pseudogenziation analysis (Fig. 4), we can pinpoint candidates species that might have been domesticated, in particular, *L. delbrueckii* subsp. *bulgaricus* that has previously been associated with a distinct evolutionary history (58) and genomic repertoire (34) tightly linked to yogurt production.

### Perspective

A key question in domestication research asks whether, for a given domesticate, there was a single domestication event that was restricted to a particular geographic area that spread across regions or if there were multiple independent domestication events (12). For cheese starter bacteria, this remains unknown. In the case of crop plants and animals, the most likely scenario seems to depend on the domesticate and the continent (1). The origin of *S. cerevisiae* is likely Asian, but it is still unclear whether the domestication happened initially in Asia and the domesticate then spread or whether the wild ancestor spread first and then got domesticated several times in different places (19,59), reviewed in (24). In the case of cheese starter bacteria, data from across a larger diversity of continents and traditional cheese varieties will be key to differentiate between a single or a multiple-origin scenario. In addition, the data we present here do not encompass closely related “wild” strains that might share a very recent common ancestor with cheese starter culture strains. Discovering and characterising these wild relatives will be instrumental to reconstructing the evolutionary history of cheese starter culture and testing the single vs. multiple origin scenarios as well as the degree of hybridisation between domesticated and wild strains. Previous studies have already suggested that these species likely do not have an original niche in the human gut (46,60–62) potentially having a plant origin. However, we hypothesize here that the ancestral niche of cheese starter bacteria might be the gut of milk-feeding (i.e. juvenile) dairy animals that have originally been used as a source for rennet (63).

## Methods

### Community sampling

The cheese starter cultures were continuously passaged at Agroscope (Liebefeld, Switzerland) as undefined starter cultures. The process of maintaining an undefined starter culture is explained extensively in (35). In short, a stock culture is aliquoted into several samples which are used on a weekly basis to fulfil the demand of cheese makers. The initial stock is maintained as a freeze-dried ampule at 4°C and is only passaged and aliquoted if necessary. The number of passages is irregular and not documented. In general, the stock cultures were passaged more frequently from 1970 to approximately 2000, as frequently as every month since we have numerous samples dated one month apart. Today, the frequency of passaging is reduced to a minimum which can be as rare as every 5 years. The historic samples were stored at 4°C throughout the years and DNA isolation was done collectively in 2019. The DNA isolation was done as previously described (64). In short, the DNA was isolated with the EZ1 DNA Tissue kit on the BioRobot EZ1 robot (Qiagen, Hombrechtikon, Switzerland).

### Strain isolation and bacterial counts

Bacterial strains were isolated as previously explained in Somerville et al. 2022 (35). In short, all 11 cheese starter cultures were plated on two selective media SPY9.3 (65) and MR11 (MRS adjusted to pH 5.4 according to ISO7889) for *S. thermophilus* and *L. delbrueckii,* respectively. Ninety-six colonies per species were randomly picked and cultured in liquid media for 24 h at 37 °C. For genotyping, DNA from 100 μL of culture was extracted using the EtNa DNA isolation method (66) and mini-satellite PCR for strain identification of *S. thermophilus* and *L. delbrueckii* was done as described previously (35). Colony-forming units (CFU/ml) were determined by serial dilution and plate counting with an Eddy jet spiral plater and SphereFlash Automatic Colony Counter (both from IUL, Barcelona, Spain).

### Metagenome and genome sample preparation and sequencing

Ninety-eight samples including historic freeze-dried ampules, present working stocks and cheese starter cultures were prepared for shotgun metagenome sequencing. The DNA was isolated as previously explained (35), and Nextera flex libraries were prepared and subjected to HiSeq4000 150PE (Illumina) sequencing at the Genomic Technologies Facility in Lausanne, Switzerland. Further, a subset of genomic samples was sequenced on a minION (Nanopore) with a rapid barcoding kit.

### Raw read analysis

The raw reads for both metagenomic and genomic samples were handled similarly. All adaptors and barcodes were removed with trimgalore (67). Reads mapping to the cow genome from the milk were removed with KneadData (https://bitbucket.org/biobakery/kneaddata). The reads were mapped with bwa mem (68). The genomes and metagenomes were assembled with SPAdes (69) for short reads and Flye for long reads (70). Extensive genome polishing was done with Racon (71) and freebayes (72).

### Genome analysis

The genome assemblies were submitted to NCBI and annotated with PGAP (73). Additionally, eggnog mapper was used to identify annotations of pseudogenes (74). The pairwise ANI values were calculated with fastANI (75). Additional genomes and their respective PGAP of the three focal species were downloaded from NCBI Refseq (12.01.2020). Only completely assembled genomes were used as we have previously observed a substantial number of genes (and pseudogenes) not being assembled with Illumina only assemblies due to the repetitive nature of the tranposase rich genomes (57). Orthofinder was used to identify single copy core genes within the species (76).

### Metagenome analysis

The metagenomic raw reads were profiled for species abundance with mOTU2 (41). Additionally, the metagenomic reads were mapped against reference genomes of the three focal species with bwa mem (77). We performed a single nucleotide analysis (SNV) with freebayes-parallel on the genomes and metagenomes in comparison to random reference genomes (72). The observed SNVs were filtered with vcftools (78) and SNPeffect (79) to include only SNVs with a minimum allele frequency of 0.05, a read coverage of 5 and in single-copy-genes (identified with orthofinder). The metagenomic SNVs were compared to the previously identified SNVs from the reference genomes. Sub-species clade frequency was calculated by averaging the allele frequency of all sub-species clade-specific SNVs (for details, see script).

### Propagation experiment and measurements

For the propagation experiment we created a pooled starting sample consisting of one random isolate of *L. delbrueckii* and one for *S.thermophilus* per starter culture (in total 22 isolates) (see selection in Supp. table 1). The starting sample was propagated in five replicates as described in Somerville et al. 2022 (35). In short, the samples were propagated to simulate the production of cheese starter cultures. We conducted two passages per week. On the first day, 100 ul of freeze-dried sample was inoculated into 10 ml autoclaved organic milk media (BM) and incubated for 18 h at 37 °C. For the second passage on the next day, the pre-culture was inoculated into 10 ml autoclaved BM and incubated for another 18 h at 37 °C. For the final step, 100 ul of the incubated samples were transferred into a freeze-dry ampule and stored at −30 °C for at least 1 h. Thereafter, the samples were freeze-dried for 7 h until dry. For pH measurements, we used the hydroplate system (PreSens, Germany). The pH was normalized with pH standards pH 4 and 7. The measurements were done in four replicates for 30 h at 37 °C.

### Species-level lactic acid bacteria phylogeny

All representative genomes from lactic acid bacteria species were downloaded from NCBI Refseq. Moreover, from the genus *Streptococcus* and *Lactobacillus*, all genomes deposited on NCBI before 12.01.2020 were included. The annotations were screened for Galactosidase (lacZ and lacG). The phylogeny was reconstructed by concatenating the 16S, 23S and 5S rRNA sequence of the genomes, making a multi-sequence-alignment file with mafft-linsi (80) and calculating the phylogeny with RaxML and the “GTRCAT” model (81). The plot was created in R with ggtree (82).

### Strain-level dated whole genome phylogeny

The preliminary species tree was calculated with OrthoFinder (76). Thereof we back-translated the single copy core genes into the nucleotide space and created a core genome species tree and gene-trees as described in (83) using MAFFT (84) and RAxML with 100 bootstrap rounds and the GTRCAT (81). Moreover, by including the gene-tree and the species tree we calculated the clonal species tree with ClonalFrameML (85). Thereof a dated phylogeny was calculated with BactDating and 10⁷ Bayes repetitions using our historical samples as time calibration points (49). From the subsequent phylogeny, we predicted sub-species clades for our genome isolates with poppunk (44). In order to quantify the presence of sub-species clades in the metagenomes, we identified all sub-species clade specific SNVs in the core genes. The dated phylogeny and the information of isolation source (Supp. table 1) were used to reconstruction ancestral habitat reconstruction with the ace function from the ape package (86).

### Transcriptome analysis

Samples for transcriptome analysis were collected throughout the first 24 hours of a regular gruyere-type cheese making process at the cheese pilot-plant at Agroscope (Liebefeld, Switzerland). The samples were immediately stored in liquid nitrogen and the RNA extraction was carried out with the Qiagen EZ1 extraction robot and the RNA Tissue kit. Illumina libraries were prepared with the TruSeq Str-RNA Zero and sequencing was performed on a 150 PE HiSeq 4000 (Illumina) at the Genomic Technologies Facility in Lausanne, Switzerland. The Illumina sequences were cleaned with trimgalore (67) and sortmeRNA (87). Mapping of reads to isolate reference genomes was performed with bwa mem (77) and gene counting with HT-seq (88). Further, sample and gene normalization was performed with DESeq2 (89).

### Phenotypic assays

The carbohydrate utilization profiles were determined using Biolog™ phenotypic microarrays plates PM01 and PM02A. Samples were prepared according to the manufacturer’s protocols. Plates were incubated in Omnilog ™ for 48 h at 37 °C. The equipment records the contrast difference every 15 minutes to generate growth curves. This data was evaluated using Biologs data acquisition software ™ and the opm package in R (90). The titratable acidity of the cultures was determined as follows: 10 mL reconstituted skim milk was inoculated using 0.1 ‰ of the corresponding culture and incubated for 18 h at 38 °C. 1 drop of phenolphthalein was added to the sample and then titrated with 0.1 M NaOH till a visible colour change was detected. The recorded volume of 0.1 M NaOH in mL was multiplied by 10 and rounded to the whole number resulting in the determined °Th (or clark degree). Total lactate (D- and L-Lactate) was analysed enzymatically according to the instruction protocol of the kit manufacturer (Boehringer, Manheim, Germany) using an automated spectrophotometric analyser (Gallery, Thermo, Switzerland). The proportion of L-lactic acid to total lactic acid was calculated as a percentage. This method has previously been published (91). Linear mixed effects model were fitted into the phenotypic data with the lmer function in the lme4 package (92).

### Statistical analysis and plotting

If not mentioned otherwise, the analyses, statistics and plotting was done in R (R Core Team, 2020) using ggplot2 (93). The wilcox.test and chisq_test from the rstatix package were used.

## Data and code availability

Whole genome sequencing of illumina only genomes (BioProject: PRJNA717134) of ONT and Illumina genomes (BioProject: PRJNA717134 and PRJNA1083966). The shotgun metagenomic reads are deposited on SRA (BioProject: PRJNA1048529). All dataframes, figures and output are available on Zenodo (10.5281/zenodo.10783581). The complete code is available at: (https://github.com/Freevini/Starter-Culture-diversity).

## Author Contributions

VS: Conceptualization, formal analysis, Funding Acquisition, Visualization, Writing – Review & Editing, UvA, FM, PE: Funding Acquisition, Writing, Conceptualization – Review & Editing. NT, AR, LB, MH, HB, NS, RSS: Methodology

## Acknowledgment

We thank Xavier Didelot for the useful advice for calculating the molecular clock. Moreover, I thank Herwig Bachmann and Bas Teusink for the many discussions on these topics. Moreover, I want to thank Julien Marquis and Johann Weber from the Genomic Technologies Facility in Lausanne and Alban Ramette from the IFIK for the sequencing. FM thanks Uni. of Lausanne, NCCR Microbiome, SNF and Antoine Guisan. VS thanks the Doc.Mobility funding from the SNF (P1LAP3_199476).

