## Supplemental_file for "Genomic and phenotypic imprints of microbial domestication on cheese starter cultures"

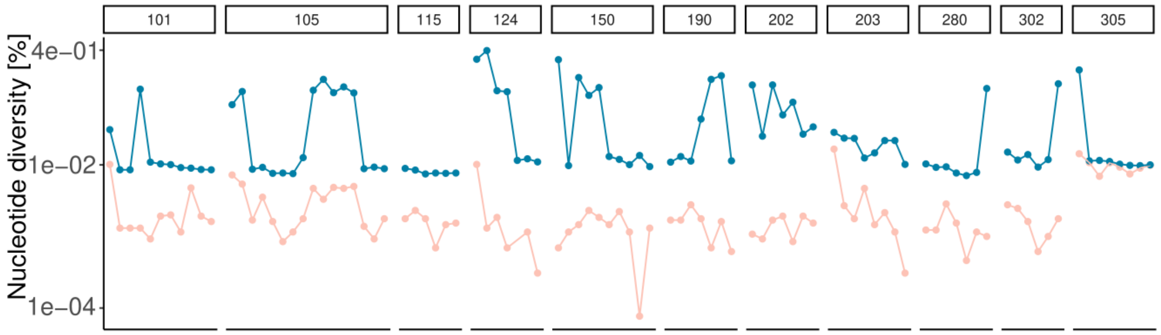


**Supplemental Figure 1. Percentage of polymorphic sites over the 98 metagenomic samples for the two focal species *S. thermophilus* (blue) and *L. delbrueckii* subsp. *lactis* (pink) (hereafter *L.delbrueckii*).** Samples (X-axis) are grouped by cheese starter culture and ordered from most ancient to most recent (see Figure 1 of main text).

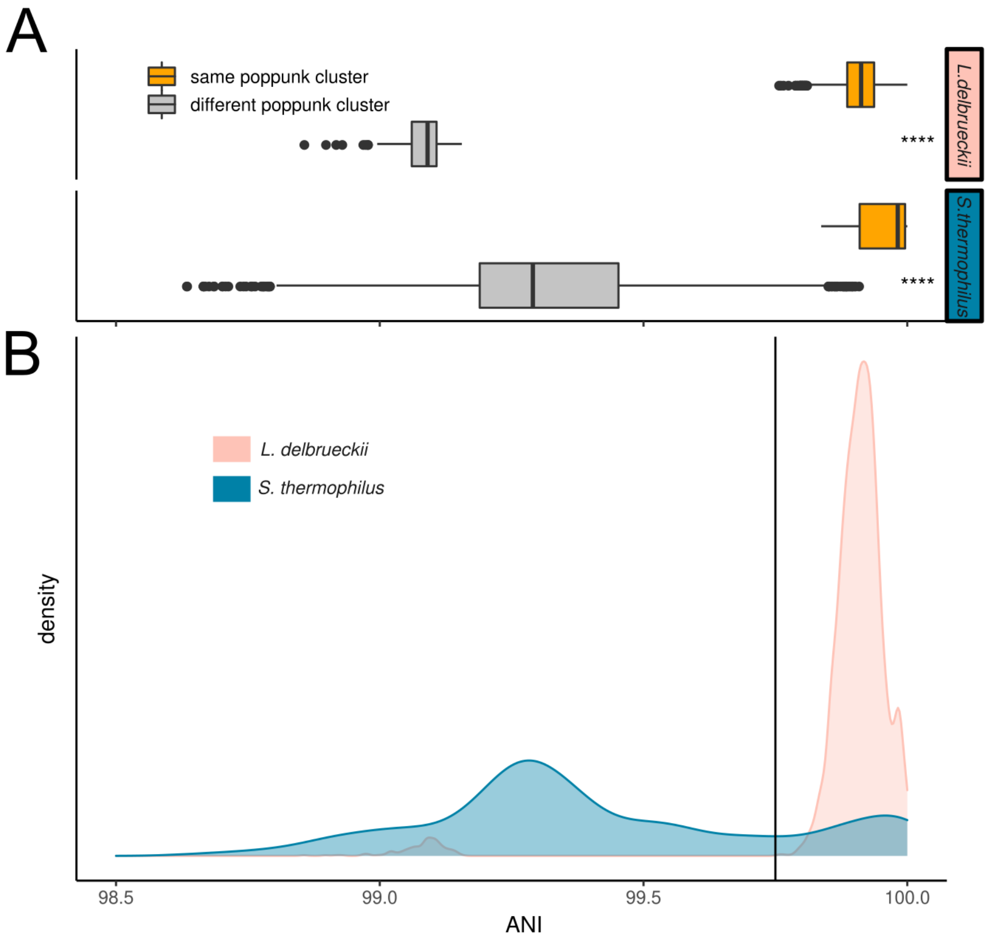


**Supplemental Figure 2: The average nucleotide identity (ANI) between and within PopPUNK clusters**. These clusters are then defined as sub-species clades in the main text. The data for *L. delbrueckii* and *S. thermophilus* is illustrated as A) boxplot and correspondingly as B) density distribution of pairwise ANI values. For *L. helveticus* we only identified 1 isolate.

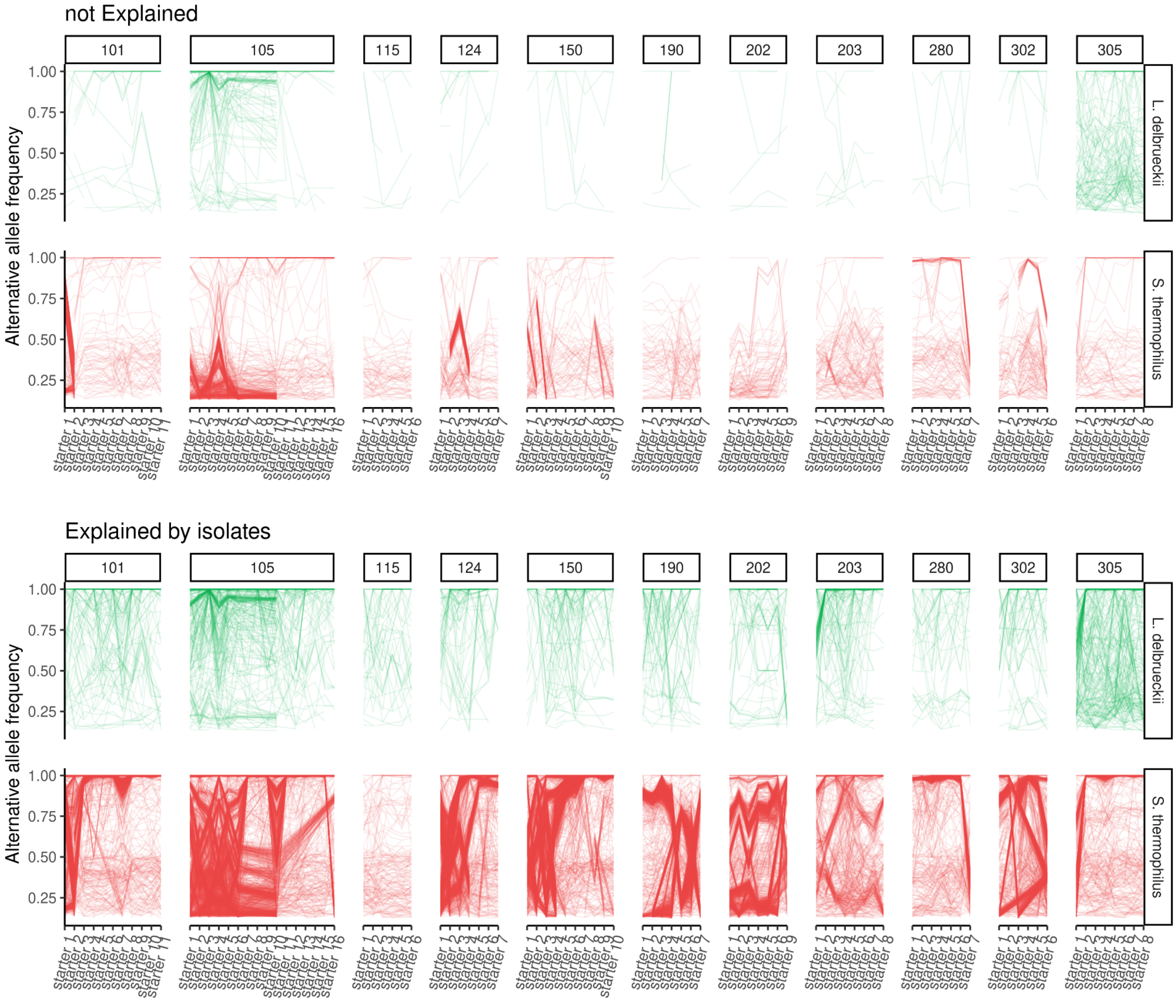


**Supplemental Figure 3: Metagenomic SNPs recovered or not by reference genomes.** The SNPs that are not explained (top) and are explained (bottom) by the isolates for both focal species. Here, “explained” means that a given SNP from the metagenomes is recovered in the reference genomes. The individual lines represent the same SNPs and the y-axis illustrates the alternative allele frequency of the corresponding SNPs. X axis corresponds to the samples, grouped by the cheese starter culture.

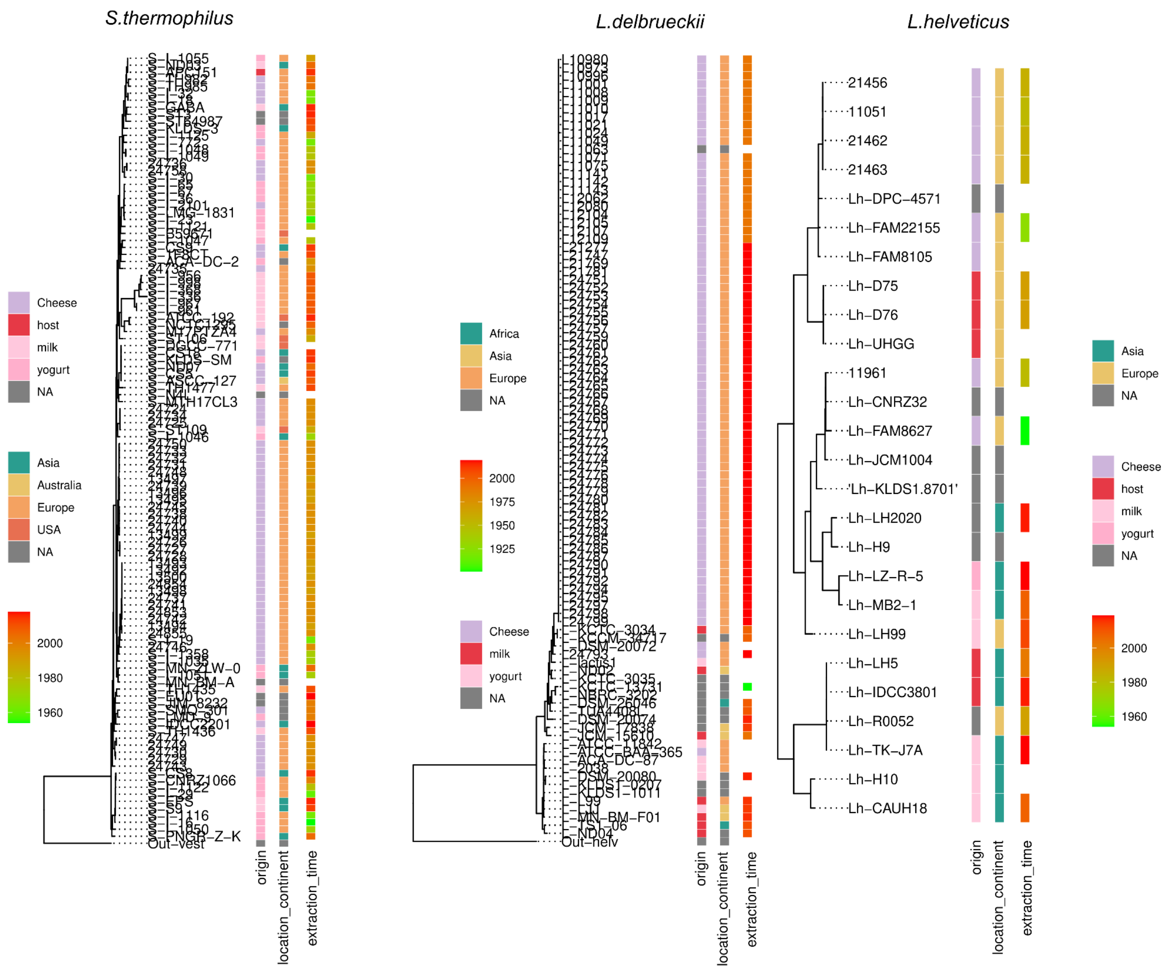


**Supplemental Figure 4: Phylogenies of the three focal species.** This figure completes the main text figure 3 by including other information on the strains including 1) isolation origin, 2) isolation geographic location and 3) isolation time.

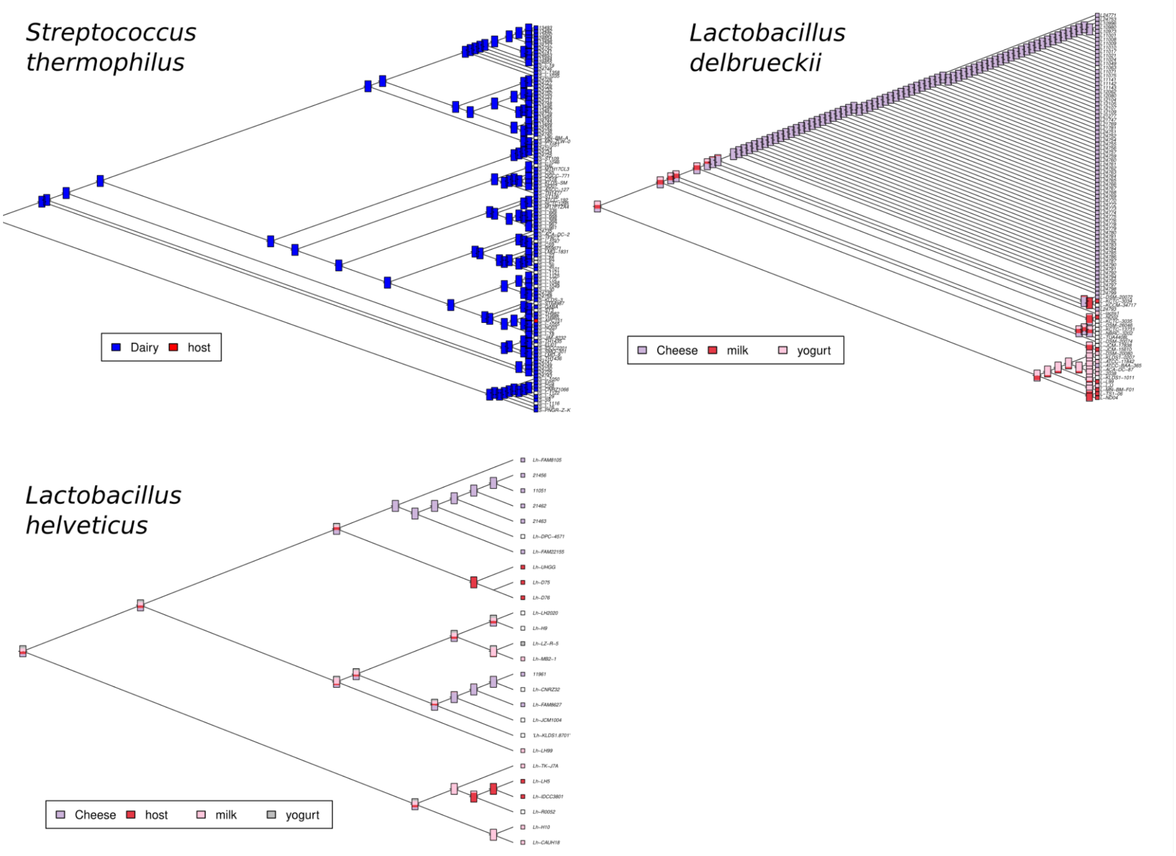


**Supplemental Figure 5: Phylogenies and ancestral habitat reconstruction of the three focal species.** The figures depict the inferred ancestral habitats for the strains of the three focal species using ace from the vegan package.

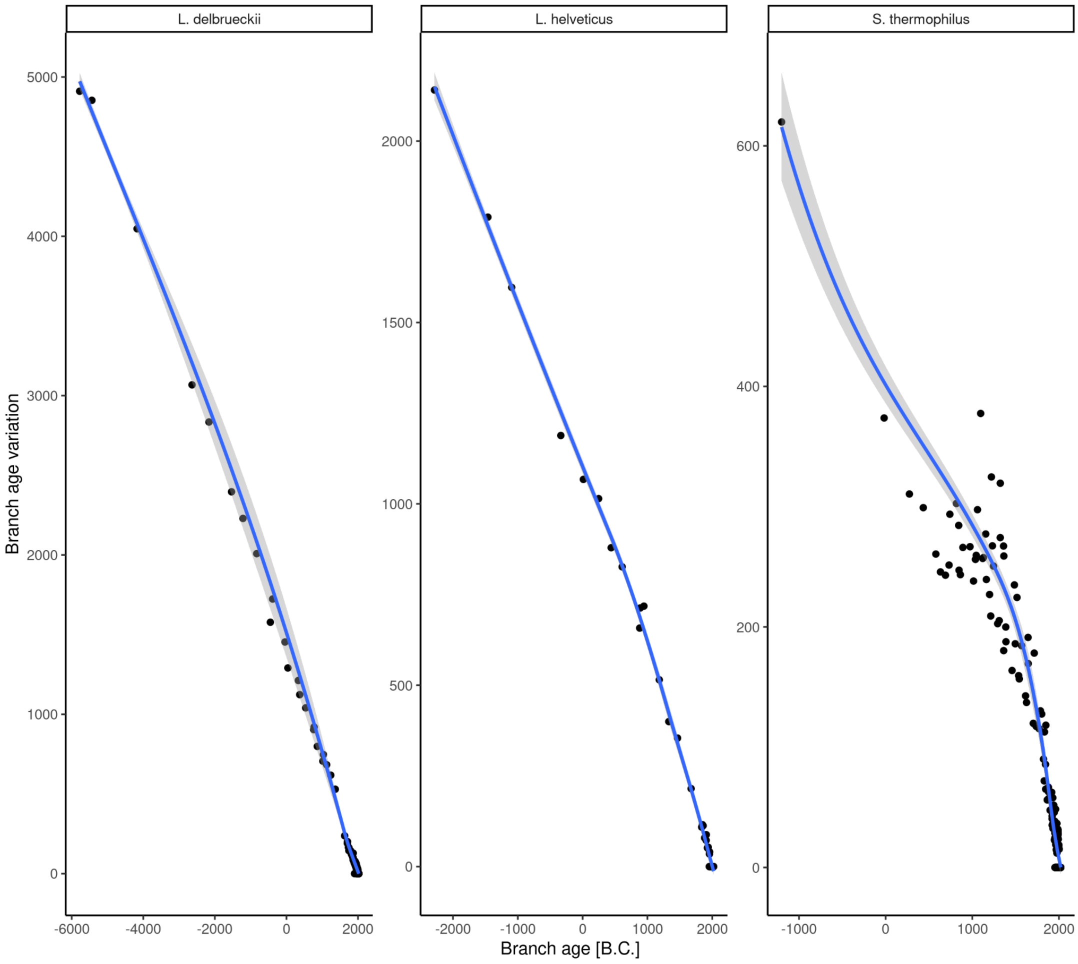


**Supplemental Figure 6:** The dated phylogeny branch age and variation of the branch age. This illustrates that the older the branches the higher the associated variation (or error) is. However, also when including the branch age variation, the crown age (oldest branch) is always after 0 B.C.

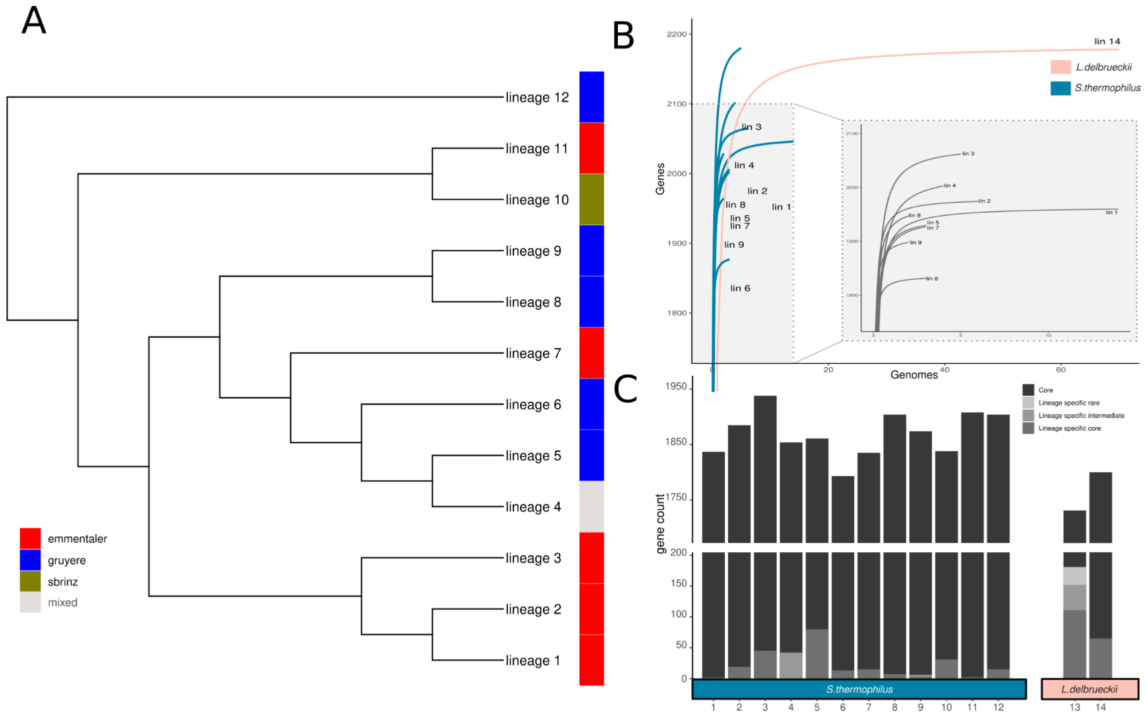


**Supplemental Figure 7: Phylogeny and gene content of reference genomes.** The A) core-gene phylogeny, B) pan-genome and C) gene content of the different sub-species clades of *S. thermophilus* and *L. delbrueckii.*

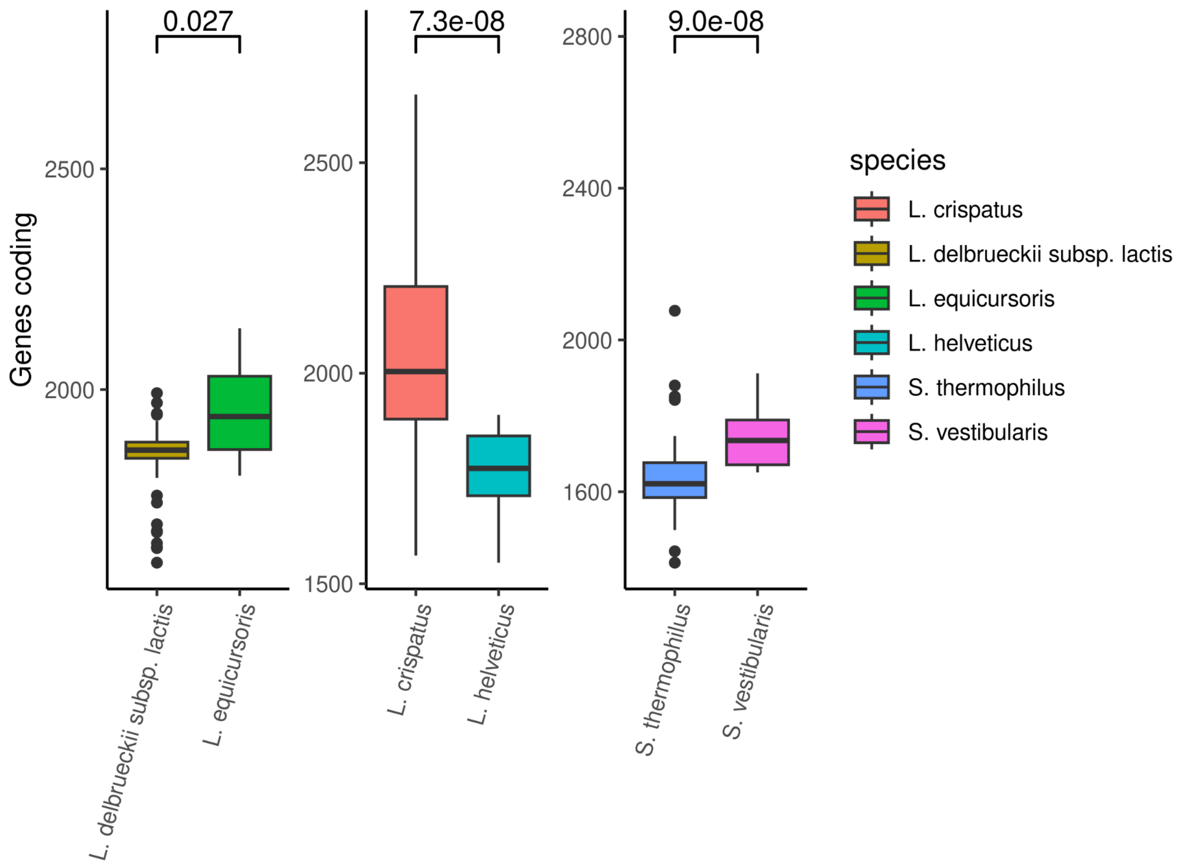


**Supplemental Figure 8: The number of coding genes in the cheese starter culture species and their closely related (sister) species**. Statistical significance is based on a Wilcoxon test and is indicated above the boxes.

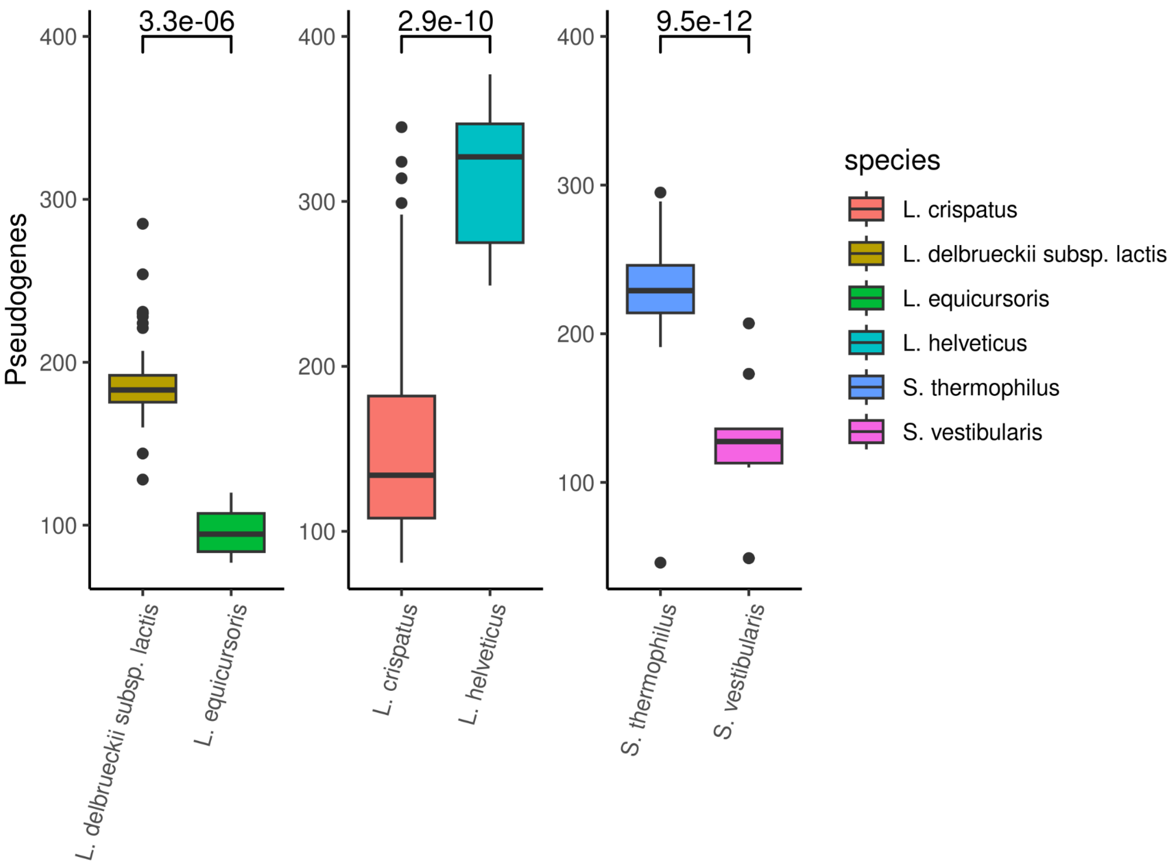


**Supplemental Figure 9: The number of pseudogenes in the cheese starter culture species and their closely related (sister) species**. Statistical significance is based on a Wilcoxon test and is indicated above the boxes.

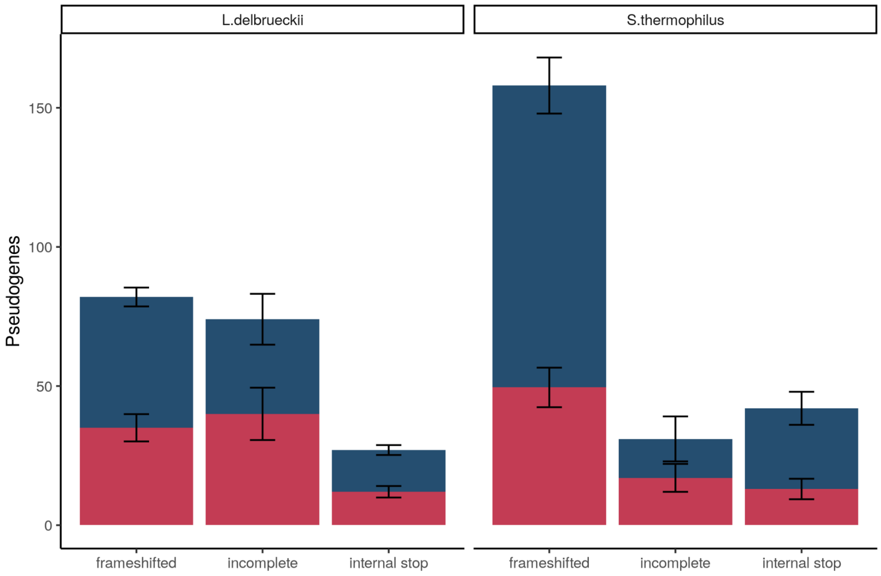


**Supplemental Figure 10: Mechanisms of pseudogenization**. The fraction and number of pseudogenes in the different species and strains that have been pseudogenized by transposons (RED) or other mechanisms (BLUE).

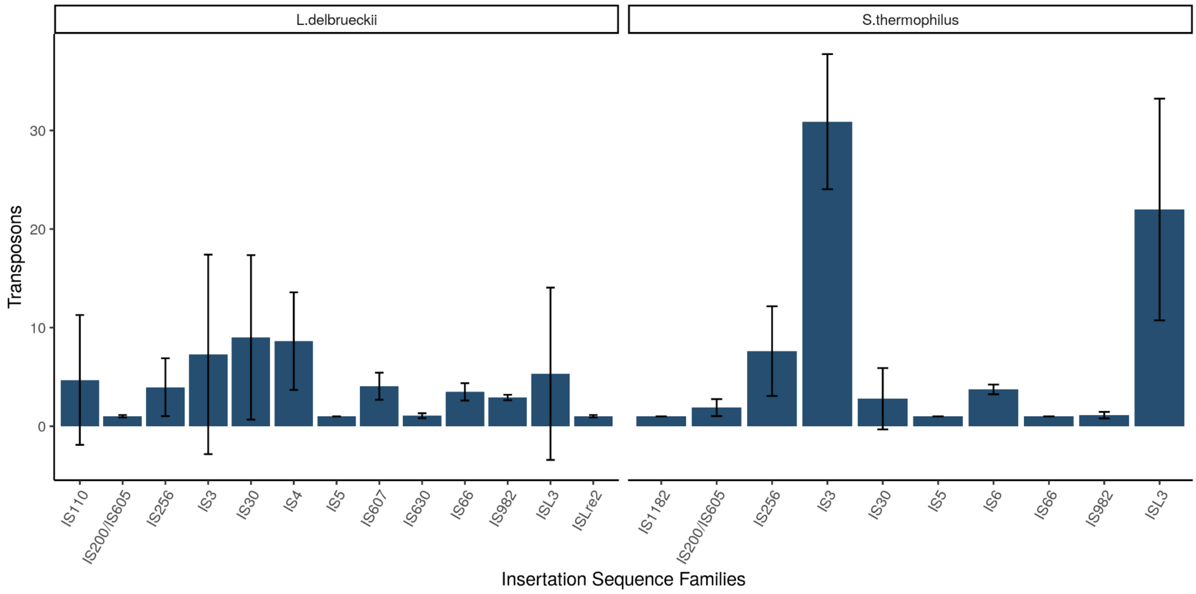


**Supplemental Figure 11: The number and variety of different insertion sequence families in the different species.** For each genome in each species, the number of transposons (y-axis) in different families (x-axis) is reported. The error bars illustrate standard deviations in the different strains.

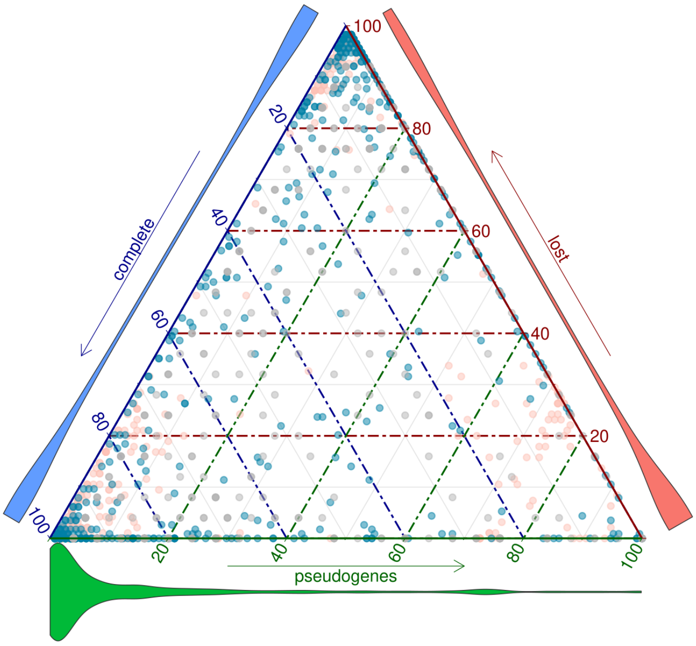


**Supplemental Figure 12: The fraction of pseudogenes, complete genes, or lost genes in orthologous gene families (OGs) that contain at least one pseudogene.** The colours of the points correspond to the species indicated in the legend in main figure 1. The violin plots at the edges indicate the distribution of the corresponding values (complete, lost or pseudogenized).

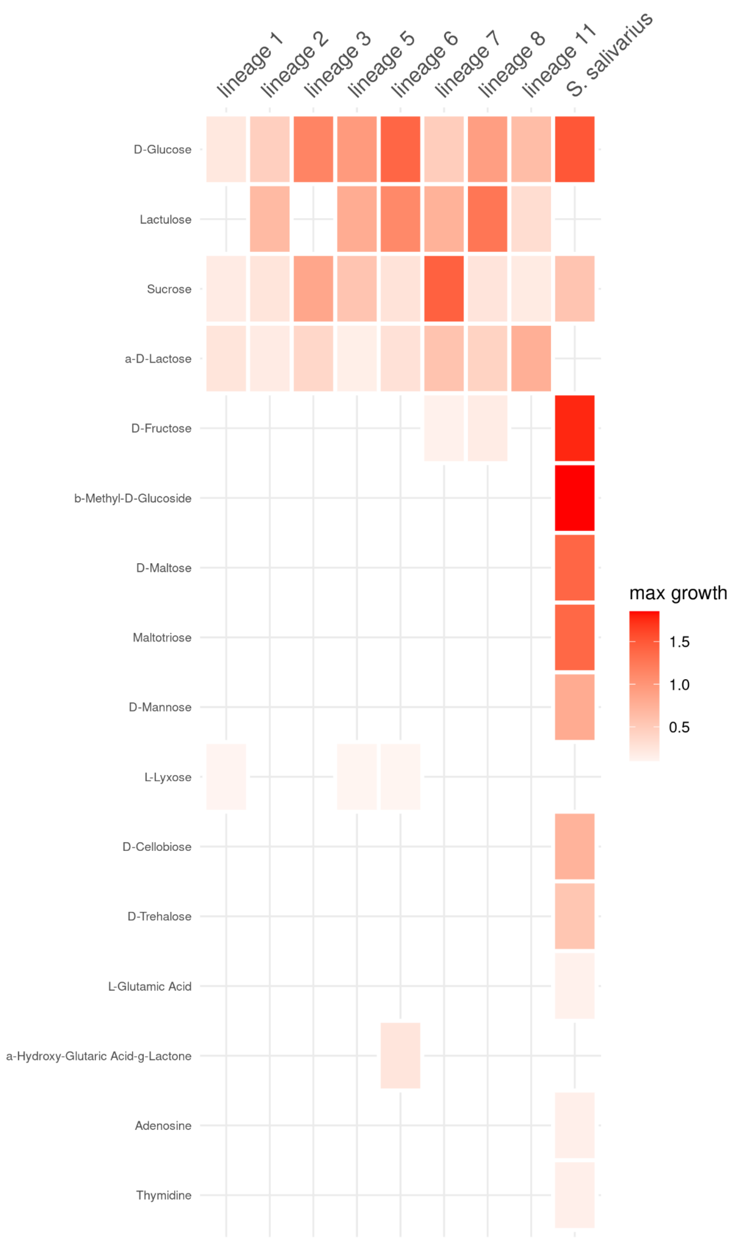


**Supplemental Figure 13: Growth assays of *Streptococcus* on various carbon sources.** The max growth rate (umax) of six *S. thermophilus* strains compared to *S. salviarius* on the different substrates of the BiologTM
100 plates. Only the substrates that had growth in any of the strains are illustrated.

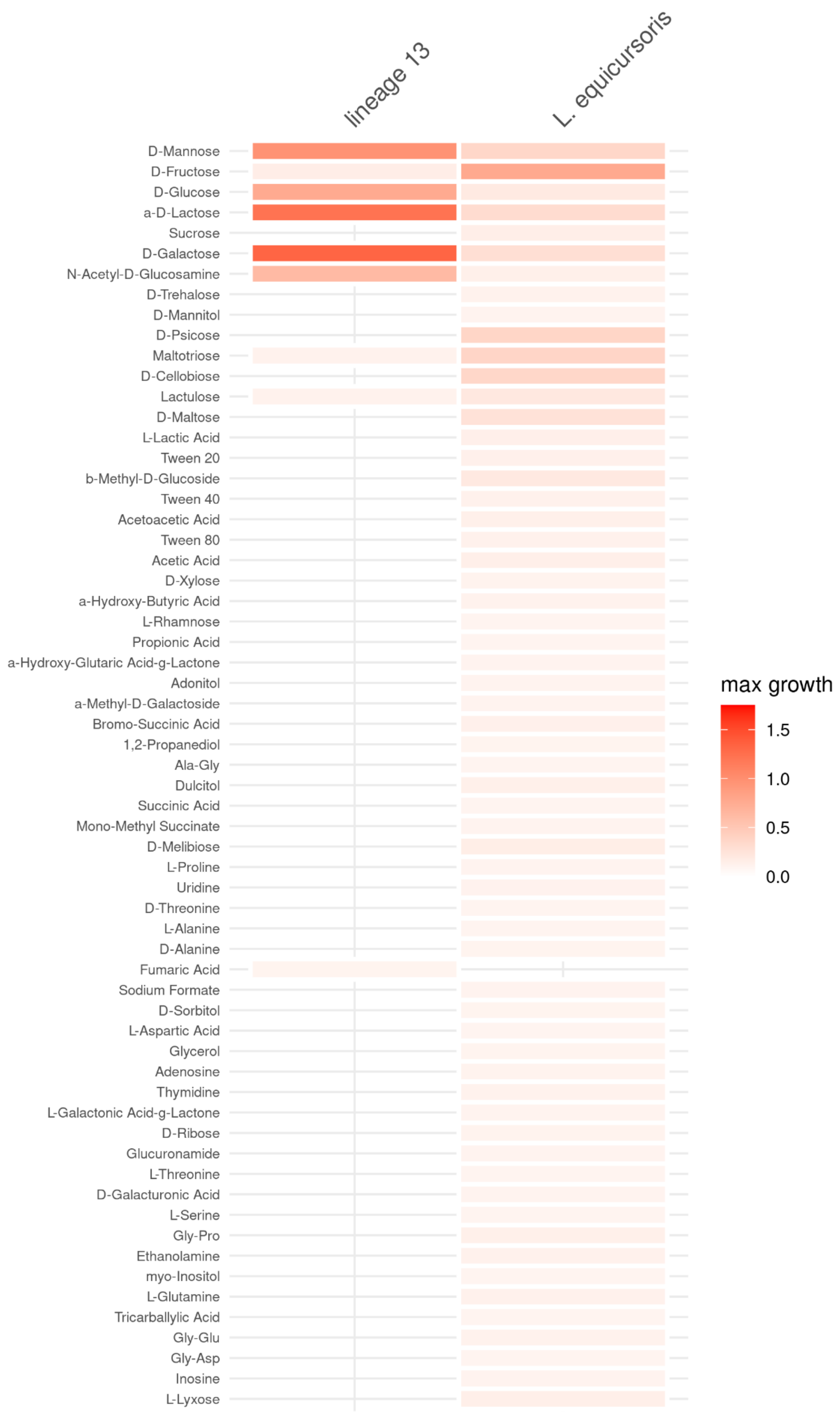


**Supplemental Figure 14: Growth assays of *Lactobacillus* on various carbon sources.** The max growth rate (umax) of the *Lactobacillus delbrueckii* strain compared to *Lactobacillus equicursoris* on the different substrates of the BiologTM plates. Only the substrates that had growth in any of the strains are illustrated.

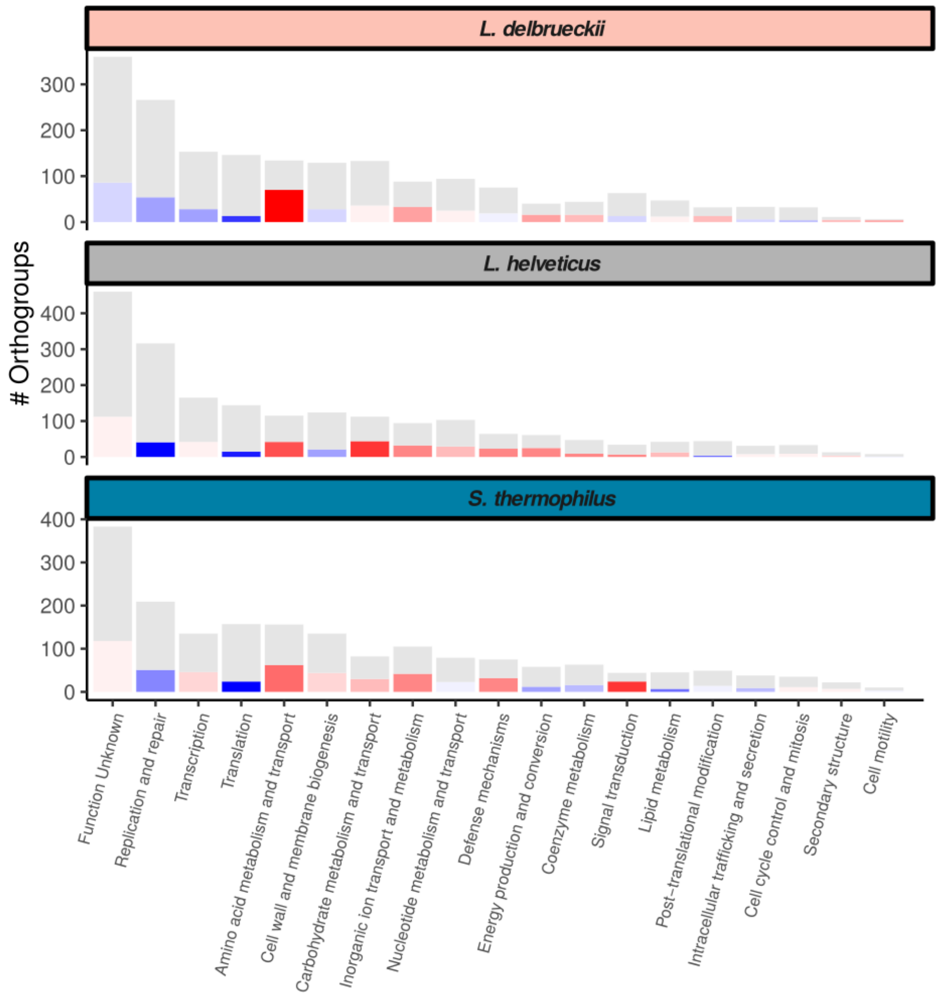


**Supplemental Figure 15: Orthologous gene family counts and proportion of pseudogenes.** The total number of orthologous gene families (OGs) (grey) and the OGs containing pseudogenes coloured red (more) or blue (less) than expected per COG category by Chi-square test. The three different species are separated by row accordingly.

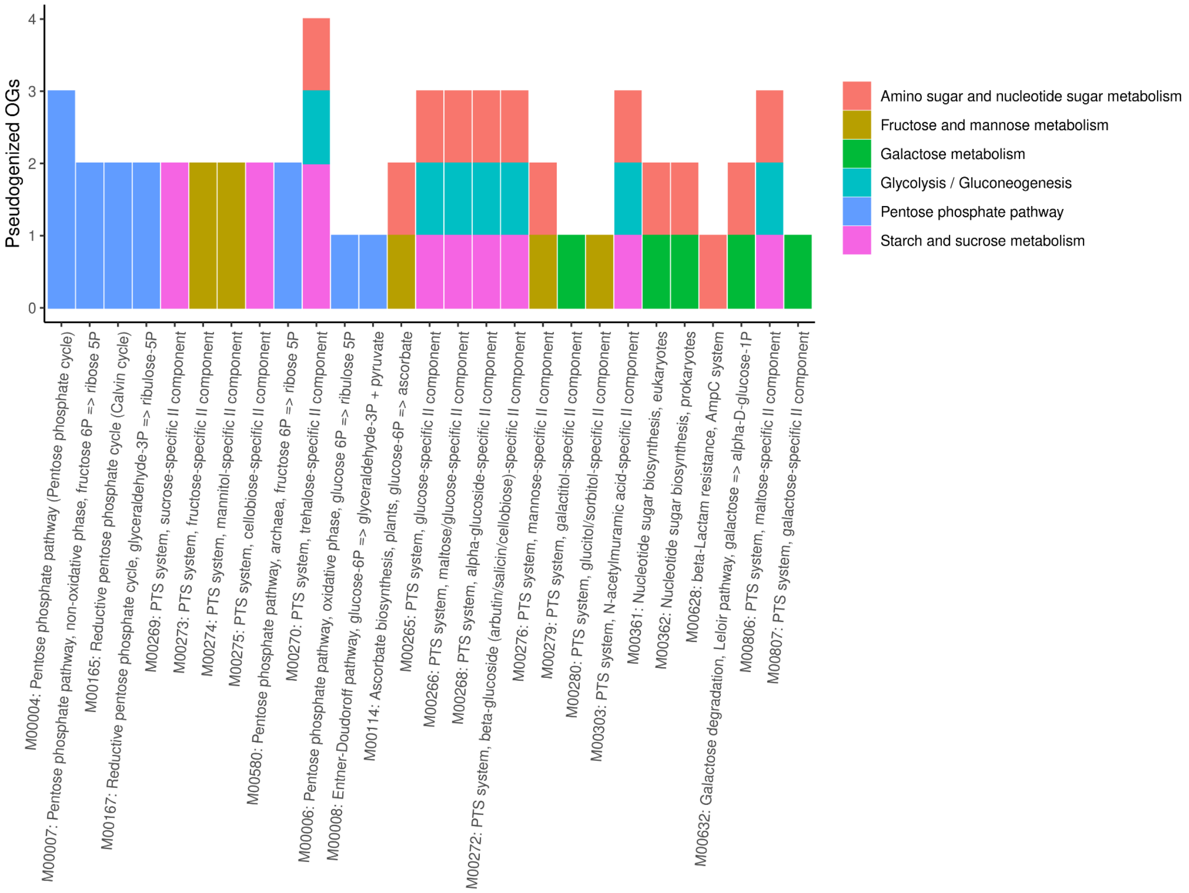


**Supplemental Figure 16: Metabolic KEGG pathway annotation of pseudogenes**. annotation of the carbohydrate genes into different KEGG metabolic modules which have been pseudogenized.

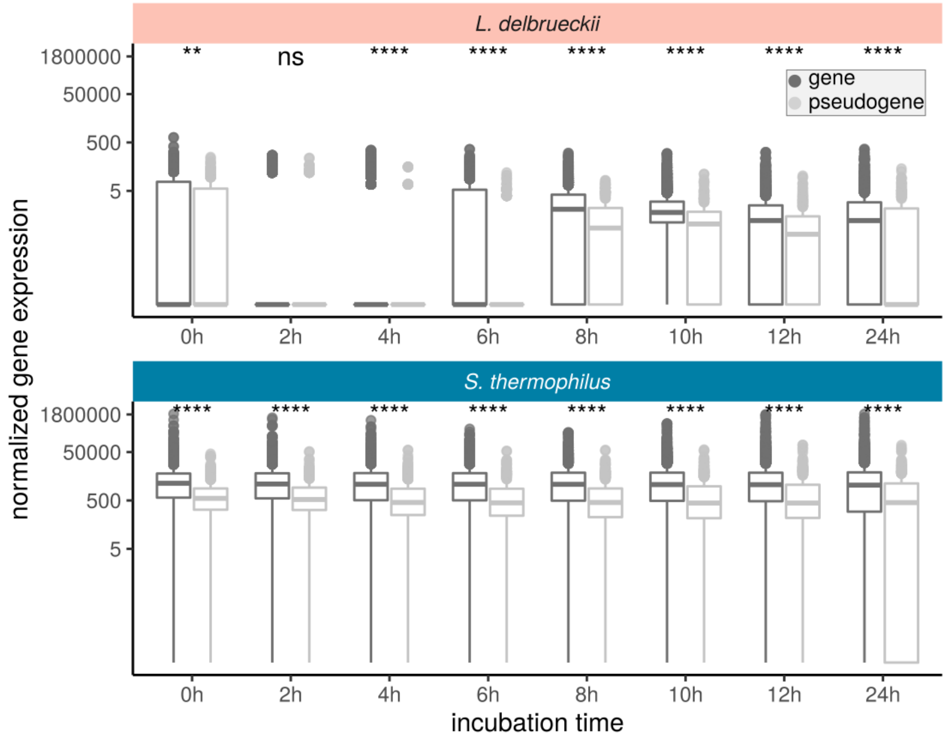


**Supplemental Figure 17: RNA-seq experiment.** The normalized gene expression of pseudogenized and non-pseudogenized genes of both species throughout the first 24h of cheese making (incubation time). The significance of the Wilcoxon test between the two groups is illustrated by stars (ns: p > 0.05, *: p <= 0.05, **: p <= 0.01. ***: p <= 0.001, ****: p <= 0.0001)
